# Pervasive differential splicing in Marek’s Disease Virus can discriminate CVI-988 vaccine strain from RB-1B virulent strain in chicken embryonic fibroblasts

**DOI:** 10.1101/723114

**Authors:** Yashar Sadigh, Abdessamad Tahiri-Alaoui, Stephen Spatz, Venugopal Nair, Paolo Ribeca

**Affiliations:** The Pirbright Institute, Ash Road, Woking, GU24 0NF, United Kingdom; Clinical BioManufacturing Facility, Jenner Institute, University of Oxford, Old Road, Headington, Oxford OX3 7JT, United Kingdom; US National Poultry Research Center, 934 College Station Road, Athens, GA 30605, United States

## Abstract

Marek’s disease is a major scourge challenging poultry health worldwide. It is caused by the highly contagious Marek’s disease virus (MDV), an alphaherpesvirus. Here we show that, similar to other members of its *Herpesviridae* family, MDV also presents a complex landscape of splicing events, most of which are uncharacterised and/or not annotated. Quite strikingly, and although the biological relevance of this fact is unknown, we found that a number of viral splicing isoforms are strain-specific despite the close sequence similarity of the strains considered, virulent RB-1B and vaccine CVI-988. We validated our findings by devising an assay that discriminates infections caused by the two strains in chicken embryonic fibroblasts based on the presence of some RNA species. To our knowledge, this study is the first ever to accomplish such a result, emphasizing how important a comprehensive knowledge of the viral transcriptome can be to understand viral pathogenesis.

**Importance:** Marek’s disease virus (MDV) causes an agro-economically important disease of chickens worldwide. Although commercial poultry are vaccinated against MDV, it is not possible to achieve sterilising immunity, and available vaccines can only protect chickens against the symptoms of the disease. Vaccinated chicken often become superinfected with virulent strains, shedding virus into the environment. The most effective MDV vaccine strain, CVI-988, shares >99% sequence identity with the prototype virulent virus strain RB-1B. Interestingly, our work shows that despite their almost identical sequences MDV strains CVI-988 and RB-1B have significantly different splicing profiles, and hence transcriptomes. We independently validated this discovery by detecting with real-time PCR some splicing isoforms expressed by MDV strain CVI-988 and absent in the transcriptome of the RB-1B strain. These results indicate that the coding potential of MDV might be much larger than previously thought, and suggest a likely underestimation of the role of the viral transcriptome in the pathogenesis and prevention of MDV.

## Introduction

Marek’s disease (MD) is a major scourge of poultry, caused by Marek’s disease virus (MDV, also known as *Gallid herpesvirus* 2, GaHV 2), a member of genus *Mardivirus* in the subfamily *Alphaherpesvirinae* of the family *Herpesviridae*. MD is characterised by paralysis, immunosuppression, and lymphoid infiltration into different tissues, including the peripheral nerves, eye, muscle and skin. Lymphoid tumours in the visceral organs can be observed as early as 3 weeks post inoculation. Control of MD has been achieved by vaccination with the live attenuated oncogenic *Gallid herpesvirus* 2 (MDV-1) strain CVI-988 (also known as Rispens) and the non-oncogenic antigenically related vaccine viruses *Gallid herpesvirus* 3 (MDV-2) strain SB-1, and *Meleagrid herpesvirus* 1 (herpesvirus of turkey, HVT) strain Fc126 (1–3). These vaccines were introduced at different periods to control the disease caused by various MDV pathotypes. Today they are used individually or more frequently in combination, since formulations including more than one vaccine can have a synergistic effect that induces stronger protection against MD (4). However, despite effectiveness of the single best vaccine (i.e. MDV strain CVI-988) in producing lifelong immunity against clinical disease and mortality, all fail to produce sterilising immunity against MDV infection. Vaccinated chickens can become infected with virulent MDV field strains – they show no obvious clinical symptoms but shed virulent virus (5). In fact there has been a continuous evolution of MDV virulence, with the emergence of hypervirulent pathotypes (6, 7). A potential role of current vaccines at driving virulence due to their inability to prevent infection and spread has been demonstrated (5). As vaccinated birds are still susceptible to superinfection by more virulent MDV subtypes, co-infection with vaccine and pathogenic strains are common in clinical materials such as poultry house dust (8).

A puzzling feature of MDV genomics is the very high sequence similarity between the virulent and vaccine strains. Despite the very different infection outcomes in vivo, the prototype virulent MDV strain RB-1B and the most effective vaccine strain, CVI-988, differ by only about 1% of their sequences. In fact the genomic differences between the RB-1B and CVI-988 strains are so limited that only one DNA-based assay (9, 10) is available to differentiate between them; the assay relies upon the detection of a few nucleotide substitutions localized to a single locus. Consequently, it would look like the differences at DNA level are not the best proxy to understand why the two strains behave so differently.

A much better vantage point might be represented by the viral transcriptome – for one thing, it has already been shown in the past that a number of herpesviruses have an unsuspectedly rich repertoire of RNAs (11–14). Detailed knowledge of the viral transcriptomes can also potentially shed a much better light on the different mechanisms taking place during virus-host interaction. In fact while our manuscript was under review, a study conducted by Betzbach and colleagues appeared in press reporting novel genes and a comprehensive annotation of MDV in primary B cells inoculated with BAC-derived MDV strains RB-1B or CVI-988 (15). Their study confirms that the community is increasingly interested in better understanding the complexity of viral transcriptomes that arise as a response to infection by closely related strains.

The work we report on here focuses on the differential expression of splice isoforms in CEF cells. Our primary goal was to establish a way of distinguishing pathogenic MDV from the CVI-988 vaccine strain by the detection of RNA transcripts differentially expressed by either strain in CEF cells. That led to the discovery of pervasive strain-dependent splicing, with markedly different splicing isoforms (spliceforms) expressed despite the fact that the two strains are closely related. To the best of our knowledge, this is the first time that such a phenomenon is reported in the literature. In the work presented here we have also directly validated in vitro, using a technique based on real-time PCR, the differential expression of three splicing isoforms that are produced only by CVI-988. That represents a solid starting point in order to be able to discriminate the infections by the two MDV strains based on the differences between their transcriptomes.

## Results

Briefly, we infected chicken embryo fibroblasts (CEFs) with two strains of MDV: virulent RB-1B and the vaccine strain CVI-988. RNA was extracted from them, and a polyA-enriched fraction was used to prepare complementary DNA (cDNA). cDNA was sequenced using Illumina technology (HiSeq 2000/2500) with a directional protocol. Reads were mapped to several MDV strains using a sensitive analysis pipeline (see Materials and Methods for more details).

The number of reads mapped with high alignment scores to the genome of CVI-988 or RB-1B, as determined by the analysis pipeline, are shown in Table 1. Raw reads have been deposited into the NCBI Sequence Read Archive under project accession number PRJNA541962.

**Table 1.** Alignment statistics for our dataset (reads have been produced using a directional sequencing protocol). To each read one or more alignments can correspond (for instance, a read might align to both terminal repeats).

| MDV strain | Reads mapping to the forward viral strand | Alignments to the forward viral strand | Reads mapping to the reverse viral strand | Alignments to the reverse viral strand |
| --- | --- | --- | --- | --- |
| CVI-988 replicate 1 | 949,845 | 1,273,630 | 639,578 | 966,599 |
| CVI-988 replicate 2 | 1,391,374 | 1,865,743 | 932,231 | 1,404,683 |
| RB-1B replicate 1 | 717,633 | 1,213,552 | 549,656 | 998,707 |
| RB-1B replicate 2 | 650,431 | 1,110,938 | 496,594 | 914,609 |

### Splicing is pervasive in MDV

In CEFs, the viral transcriptome of the MDV strains we tested consists of a complex landscape of splicing isoforms, most of which are novel and have not been reported or included in the official genome annotations.

In detail, our analysis of spliced sequencing reads (see Materials and Methods) revealed 154 introns in MDV strain RB-1B and 246 in MDV strain CVI-988 after filtering out introns with low coverage (<10 reads) and introns which are not spliced in all biological replicates (the complete list can be found in Supplementary Table 1). A small proportion of these introns, specifically those in the spliceforms of Meq, vIL18, ICP4, gC, pp38, MDV012, and LAT had already been identified and annotated (16–22) and are affirmed by this study. The localization of the introns on the MDV genome is presented in Figure 1 (see tracks 4, 6, 8 and 10), where each solid bar represents a different intron. The direction of splicing at acceptor/donor sites is indicated with a yellow arrow.

**Figure 1.**
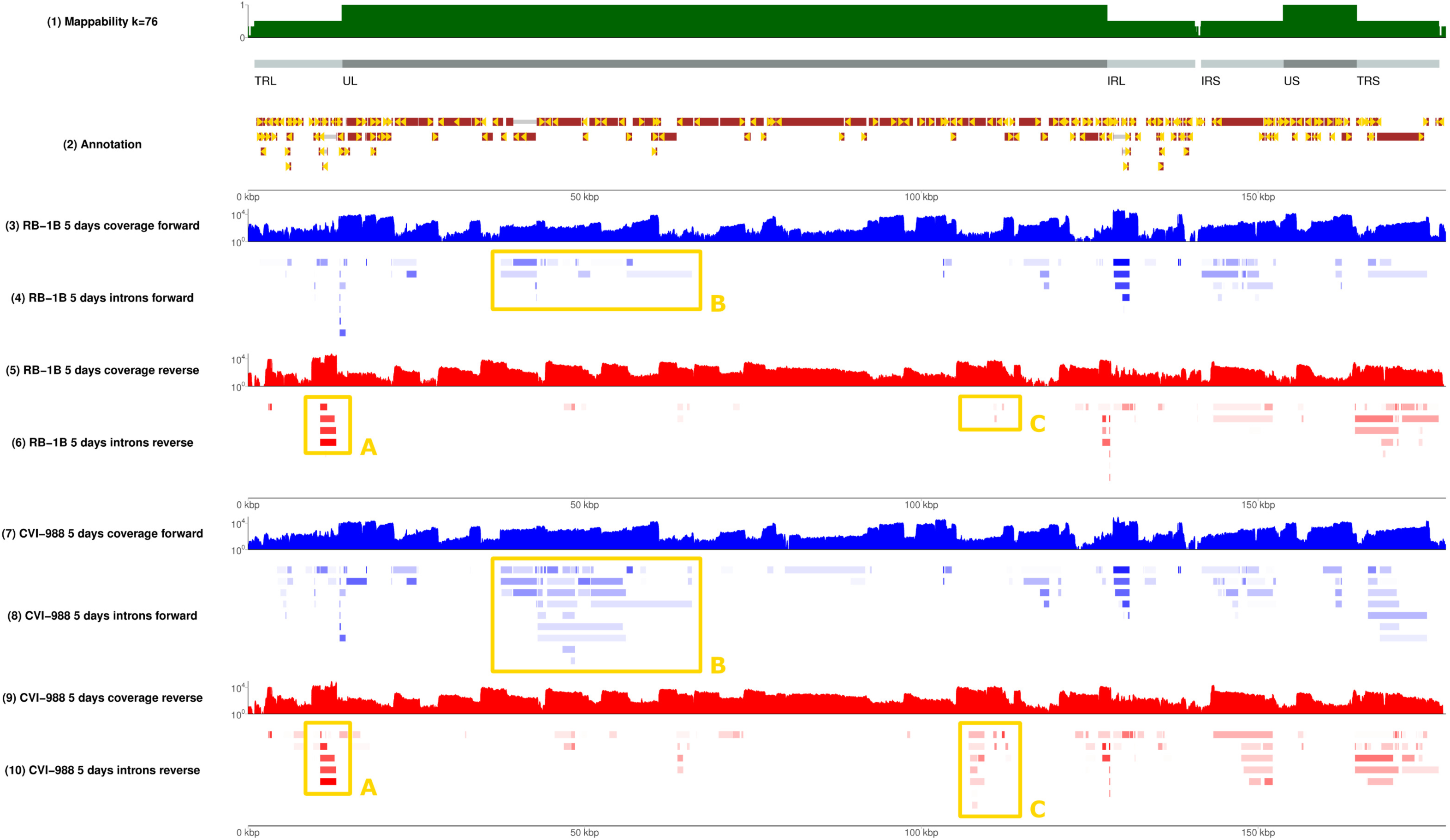
The splicing landscape of MDV infected CEFs. An overview of splicing in CEFs infected with two strains of MDV at 5 days post infection. Results are presented as a genome browser, each label on the left corresponding to a different track on the right. Track 1 (green) shows mappability – computed as in (35) – for all 76-mers in the MDV genome, which matches the read length of our RNA-sequencing experiment. Areas having mappability 1 correspond to unique regions, while areas having mappability 0.5 correspond to parts of the genome repeated twice. Track 2 (brown) shows the position of the open reading frames originally annotated by Tulman and colleagues (39–41) plus a few additional transcripts published more recently – gene nomenclature has been omitted to reduce clutter. Blue profile tracks (3 and 7) show the coverage of directional RNAseq along the forward strand in different biological conditions (infection with MDV strain RB-1B for track 3, and infection with MDV strain CVI-988 for track 7). Red tracks (5 and 9) show the corresponding coverage along the reverse strand of the virus. Tracks 4, 6, 8 and 10 show observed introns, as deduced from coverage in spliced reads. The intensity of a colour corresponds to its degree of coverage. Introns tracks are also directional (forward and reverse), and to each coverage track there corresponds an intron track (see labels). Placement of an intron in the inverted repeats (TRL vs IRL or IRS vs TRS) was done arbitrarily. There are several regions of the genome showing substantially different splicing patterns for the RB-1B and CVI-988 strains. Region A is magnified in Figure 2; region B in Figure 3; region C in Figure 4. An interactive version of the browser can be found at https://mallorn.pirbright.ac.uk/browsers/MDV-annotation/. This Figure can be reproduced in the browser by accessing the following URL: https://mallorn.pirbright.ac.uk/browsers/MDV-annotation/?loc=GaHV2_MD5%3A1..176031&tracks=Reference%2CMappability%20k%3D76%2CAnnotation%2CRB-1B%205%20days%20coverage%20forward%2CRB-1B%205%20days%20introns%2CRB-1B%205%20days%20coverage%20reverse%2CCVI-988%205%20days%20coverage%20forward%2CCVI-988%205%20days%20introns%2CCVI-988%205%20days%20coverage%20reverse&highlight=.

A number of positions appear to be alternative splice donors/acceptors, with several possible corresponding introns being selected by the splicing machinery (see Figure 2). The likelihood of such choice, which can be evaluated from RNA-sequencing data by comparing the coverage of the different splice junctions, is highly variable – it can be very similar or very different depending on the specific splice site. The distribution of splicing donors/acceptors is also not uniform across the whole genome, but rather concentrated in some specific regions, especially the inverted repeats and, surprisingly, the unique long region between 30 and 70kb (see Figure 1 and 3).

**Figure 2.**
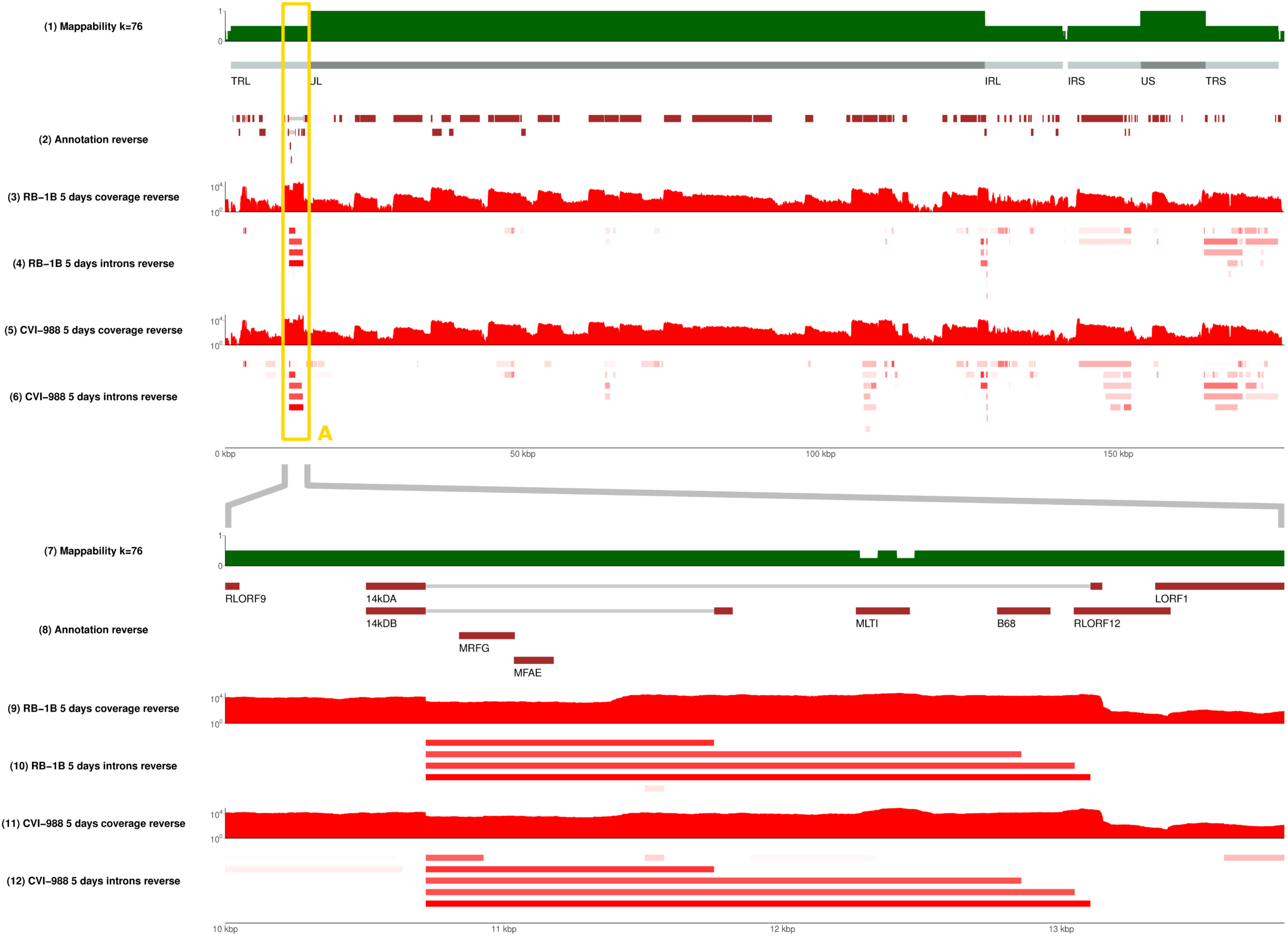
The splicing landscape at the junction between terminal repeat long and unique long regions. Alternative spliceforms across the 14KD polypeptide (pp14) gene of MDV in MDV strain RB-1B and MDV strain CVI-988 are compared. The top panel shows the whole MDV genome with the bottom panel showing a magnification of the region from 10 to 14kb (TRL/UL junction). From the RNA-sequencing signal one can see 4 main alternative spliceforms in CEFs infected by RB-1B, whereas 5 alternative spliceforms were identified in cells infected by CVI-988. As shown in track 8 (Annotation) only 2 isoforms (14 kDA and 14kDB) are present in the standard MDV annotations as defined by Hong and Coussens (42). The bottom panel of this Figure can be reproduced in the MDV genome browser by accessing the following URL: https://mallorn.pirbright.ac.uk/browsers/MDV-annotation/?loc=GaHV2_MD5%3A9554..14778&tracks=Reference%2CMappability%20k%3D76%2CAnnotation%2CRB-1B%205%20days%20coverage%20reverse%2CRB-1B%205%20days%20introns%2CCVI-988%205%20days%20coverage%20reverse%2CCVI-988%205%20days%20introns&highlight=.

**Figure 3.**
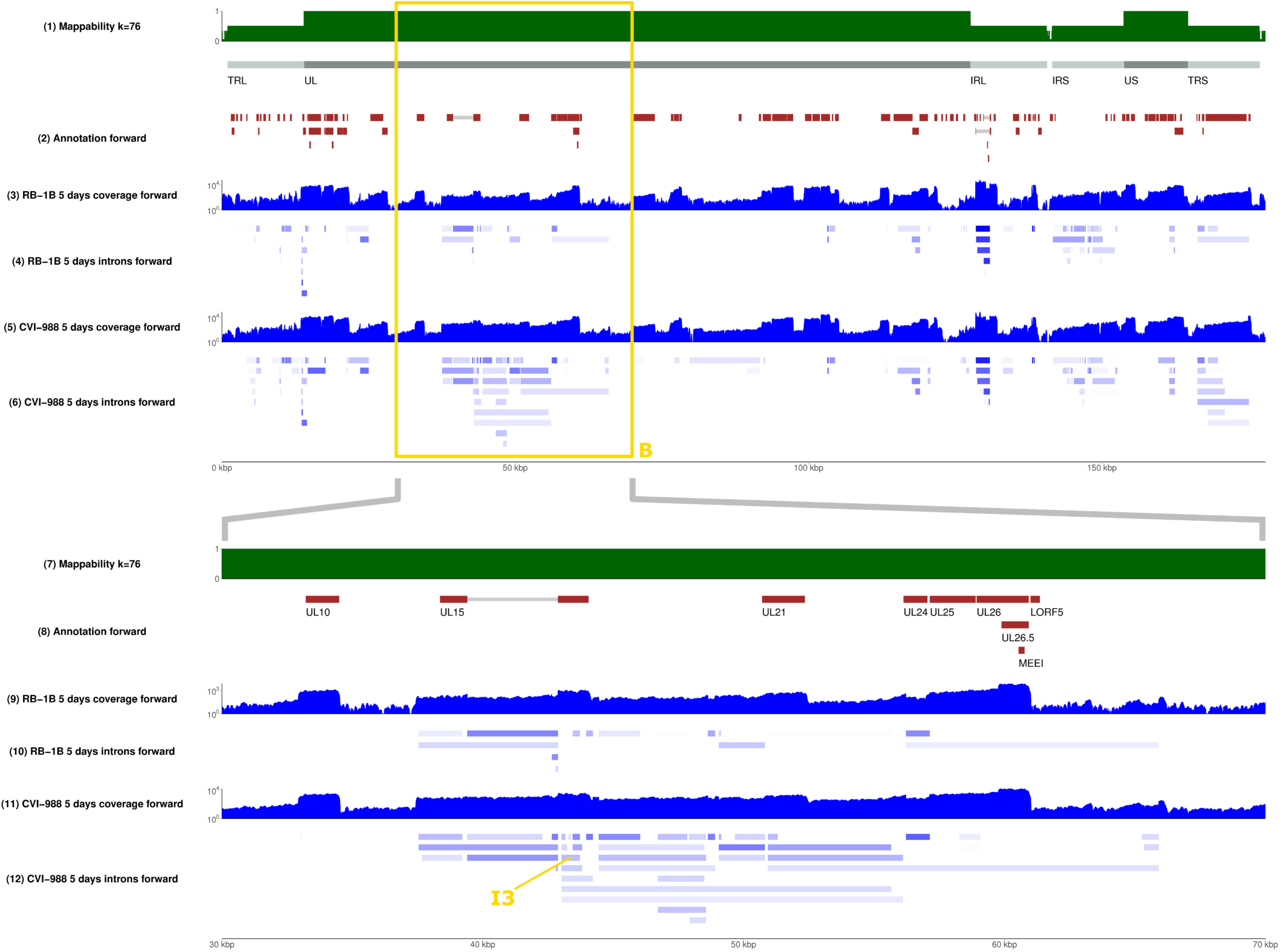
The landscape of alternative splicing in the unique long region of MDV. Splicing events on the forward strand of MDV (top panel). The region between positions 30 and 70kb is magnified in the bottom panel. Although many annotated and unannotated introns are present along this area in RB-1B during its infection of CEFs (track 9), the complexity of the splicing landscape in the same region is vastly superior in CVI-988, which shows several times more introns being actively spliced (track 11). Intron I3, which is one of the several expressed in CVI-988 and not in RB-1B, has been highlighted in the figure. The bottom panel of this Figure can be reproduced in the MDV genome browser by accessing the following URL: https://mallorn.pirbright.ac.uk/browsers/MDV-annotation/?loc=GaHV2_MD5%3A31046..52165&tracks=Reference%2CMappability%20k%3D76%2Cnnotation%2CRB-1B%205%20days%20coverage%20forward%2CRB-1B%205%20days%20introns%2CCVI-988%205%20days%20coverage%20forward%2CCVI-988%205%20days%20introns&highlight=.

An interactive version of the data, presented as a genome browser, can be publicly accessed at https://mallorn.pirbright.ac.uk/browsers/MDV-annotation/.

### Splicing is strain-specific, and occurring more frequently in MDV strain CVI-988

Many splicing isoforms appear to be produced by both strains, as expected given their similarity in nucleotide space. Despite this, one of our major findings is the identification of specific genomic regions that encode a number of isoforms which are strain specific, at least in infected CEFs. Examples are illustrated in Figure 2, where the presence of different spliceforms within the 14 kD locus is reported; in Figure 3, showing how a much greater number of spliced introns is encoded in the genome of MDV strain CVI-988 than those encoded in the genome of MDV strain RB-1B, specifically on the negative strand of the UL region (coordinates 10-70 Kb); and in Figure 4, which reveals that this is not restricted just to the negative strand. Many similar examples of varying splicing isoforms could be found throughout the MDV genomes, corroborating the observation that the splicing patterns of MDV strains RB-1B and CVI-988 are significantly different, with much higher levels of splicing occurring in CEFs infected by MDV strain CVI-988.

**Figure 4.**
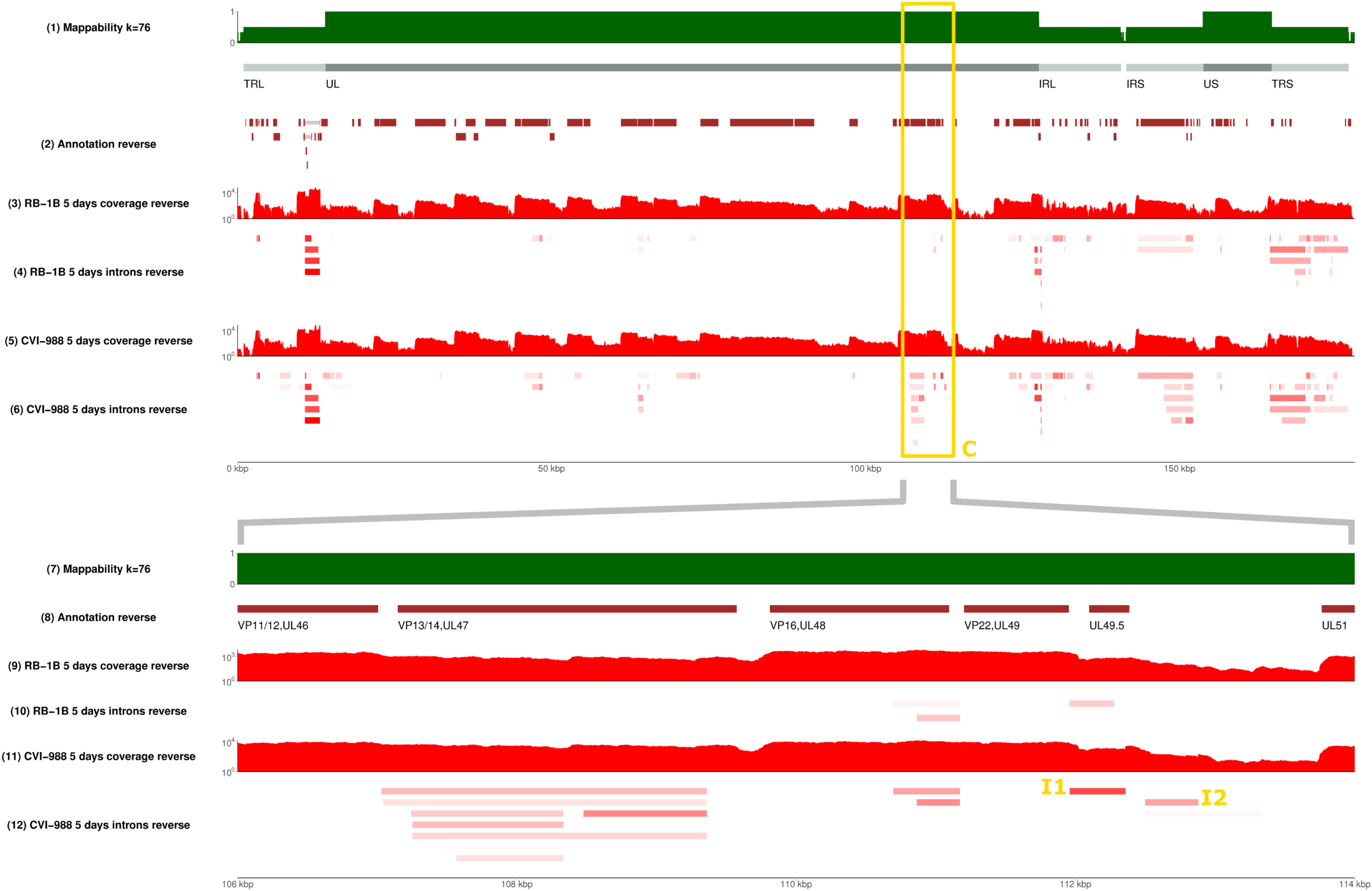
Splicing events on the reverse strand of MDV (top panel). The region between 106 and 114 kbp is magnified in the bottom panel. In this region more splicing events were observed in MDV strain CVI-988 during its infection of CEFs compared to MDV strain RB-1B. In particular one of the unique introns (named I1 in Table 3), identified in MDV strain CVI-988 between the UL49.5 and UL49 genes, was detected with a high read coverage (613 spliced reads), whereas in RB-1B the same isoform does not exist and a shorter intron can be observed. Intron I2 is also highlighted. The bottom panel of this Figure can be reproduced in the MDV genome browser by accessing the following URL: https://mallorn.pirbright.ac.uk/browsers/MDV-annotation/?loc=GaHV2_MD5%3A104493..114979&tracks=Reference%2CMappability%20k%3D76%2CAnnotation%2CRB-1B%205%20days%20coverage%20reverse%2CRB-1B%205%20days%20introns%2CCVI-988%205%20days%20coverage%20reverse%2CCVI-988%205%20days%20introns&highlight=.

### Tentative reannotation of coding transcripts

In genomic regions with many consecutive multiple splicing donors/acceptors, such as the one shown in Figure 3, the combinatorics of alternative splicing are too complex to be solved with short reads. However, it is possible to exhaustively enumerate all coding spliced transcripts compatible with the introns observed. Here, we have taken such an approach to map sequences of putative exons to all open reading frames which would translate into a protein longer than 35 amino acids (see Materials and Methods). It should be emphasised that some of such predicted coding transcripts (in particular the more complex ones involving more than two exons) may not be present at all, and confirming their presence experimentally is outside of the scope of this manuscript. However, this tentative reannotation provides a fair representation of the splicing complexity at different viral loci. Importantly, while our tentative reannotation only considers coding transcripts, many more non-coding viral transcripts might potentially be found as well.

The reannotations can be accessed, and downloaded using the feature “Save track data”, through the genome browser mentioned above. Table 2 lists the number of putative coding spliceforms annotated for a number of relevant MDV genes when CEF cells are infected by MDV strain RB-1B or MDV strain CVI-988. In Figure 5, we illustrate in more detail four cases. In panel A, we show splicing events occurring in UL49/UL49.5 after infection by MDV strains RB-1B or CVI-988 (the latter event is referred to as I1 in Table 3). Each splicing event is unique, and the nucleotide sequences of the two spliceforms differ. In addition I1, the spliceform specific to MDV strain CVI-988, is present with higher read coverage (Table 3). Panel B describes 6 new possible splicing isoforms in the ICP4 gene of MDV strain CVI-988. Additionally we observed more splicing events which could not be assigned to the ICP4 gene itself as they were detected on the opposite strand. Panel C shows two new putative splicing isoforms for pp14 occurring in both MDV strains (in addition to the two already known) and another two novel ones occurring only in CVI-988. Panel D displays 14 novel splicing events occurring at the UL52/UL53/UL54 loci in CVI-988.

**Table 2.** Enumeration of putative novel coding splicing isoforms for several MDV genes, as deduced from introns observed in CEF cells infected by MDV strains RB-1B and CVI-988.

| Genomic region | Spliceforms<br>In CVI-988 | Spliceforms<br>In RB-1B | Function |
| --- | --- | --- | --- |
| vIL8 | 5 | 5 | Viral IL8 |
| pp14A/B | 6 | 5 | 14KD lytic proteins |
| Lip | 3 | 2 | Lipase |
| UL3 | 5 | 3 | Nuclear phosphoprotein |
| UL15 | 17 | 4 | DNA packaging protein |
| UL19 | 4 | 4 | Major capsid protein |
| UL21 | 3 | 2 | Membrane protein |
| UL24 | 2 | 2 | Reactivating from latency/immune evasion |
| UL28 | 7 | 1 | DNA packaging protein. |
| UL29 | 2 | 1 | DNA binding protein |
| UL34 | 2 | 1 | Membrane phosphoprotein |
| UL38 | 2 | 1 | Capsid protein |
| UL41 | 4 | 1 | Tegument protein |
| UL44 | 6 | 5 | Glycoprotein C |
| VP11/12 | 2 | 1 | Tegument phosphoprotein |
| VP13/14 | 11 | 1 | Tegument phosphoprotein |
| VP16 | 3 | 3 | Tegument immediate early protein |
| VP22 | 2 | 2 | Tegument phosphoprotein |
| UL52/UL53/UL54 | 14 | 4 | DNA helicase/glycoprotein K/ICP27 like proteins |
| RLORF14a | 3 | 3 | 38 KD protein |
| ICP4 | 5 | 1 | IE protein |
| ICP4 area | 15 | 7 |  |
| US7 | 5 | 3 | Glycoprotein I |
| US8 | 3 | 3 | Glycoprotein E |

**Table 3.** List of introns that are (A) spliced exclusively in MDV strain CVI-988, (B) spliced exclusively in MDV strain RB-1B. To make splicing events occurring in different MDV strains comparable, all introns are listed in terms of coordinates based on the genomic sequence of MDV reference strain MD5 (NCBI accession NC_002229.3). Mapping the introns occurring in the RB-1B/CVI-988 transcriptomes to the MD5 genome was possible thanks to the extremely high sequence similarity (>99%) shared by MD5, RB-1B and CVI-988. A full table with intron coordinates with respect to all the three strains is provided as Supplementary Table 2.

| MDV strain | Strand | Position of first intron nucleotide | Position of last intron nucleotide | Gene | Read coverage | Name |
| --- | --- | --- | --- | --- | --- | --- |
| A. Introns only spliced in MDV strain CVI-988 |  |  |  |  |  |  |
| GaHV2 | - | 112359 | 111959 | VP22 gene, intergenic region | 646 | I1 |
| GaHV2 | - | 112881 | 112500 | UL50 | 58 | I2 |
| GaHV2 | + | 43021 | 43723 | UL15, terminase | 39 | I3 |
| GaHV2 | + | 43021 | 43166 | UL15, terminase | 42 |  |
| GaHV2 | + | 50934 | 51306 | UL21 | 38 |  |
| GaHV2 | - | 108334 | 107252 | UL46 / UL47 | 29 |  |
| GaHV2 | + | 43021 | 43808 | UL15, terminase | 28 |  |
| B. Introns only spliced in MDV strain RB-1B |  |  |  |  |  |  |
| GaHV2 | - | 111959 | 112277 | UL49 / UL49.5 | 19 |  |

**Figure 5.**
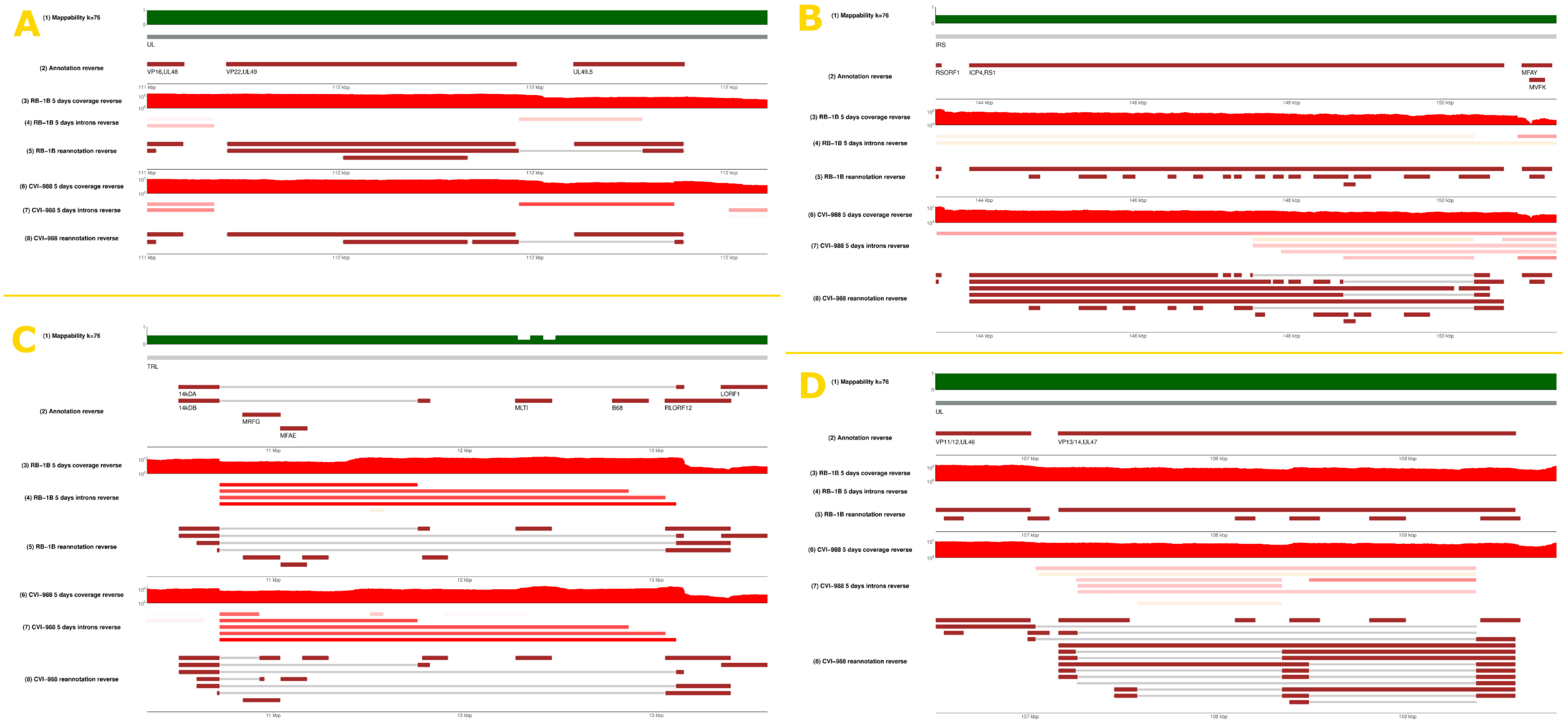
Tentative condition-dependent reannotation of splicing events on the reverse strand of MDV. In-silico reconstructed coding spliced transcripts compatible with the introns observed from Illumina short sequencing read—the coding spliceforms shown in each panel have been produced by automatic prediction methods (see Materials and Methods) but the presence of full-length transcripts has not been experimentally validated. We show the likely presence of new spliceforms for genes UL49.5/UL49 (panel A), ICP4 (panel B), PP14 (panel C), and VP13/14/UL47 (panel D), respectively. The panels of this Figure can be reproduced in the MDV genome browser by accessing the following URLs: Panel A: https://mallorn.pirbright.ac.uk/browsers/MDV-annotation/?loc=GaHV2_MD5%3A110662..112751&tracks=Reference%2CMappability%20k%3D76%2CAnnotation%2CRB-1B%205%20days%20coverage%20reverse%2CRB-1B%205%20days%20introns%2CTentative%20reannotation%20RB-1B%2CCVI-988%205%20days%20coverage%20reverse%2CCVI-988%205%20days%20introns%2CTentative%20reannotation%20CVI-988&highlight= Panel B: https://mallorn.pirbright.ac.uk/browsers/MDV-annotation/?loc=GaHV2_MD5%3A142034..152411&tracks=Reference%2CMappability%20k%3D76%2CAnnotation%2CRB-1B%205%20days%20coverage%20reverse%2CRB-1B%205%20days%20introns%2CTentative%20reannotation%20RB-1B%2CCVI-988%205%20days%20coverage%20reverse%2CCVI-988%205%20days%20introns%2CTentative%20reannotation%20CVI-988&highlight= Panel C: https://mallorn.pirbright.ac.uk/browsers/MDV-annotation/?loc=GaHV2_MD5%3A9551..14739&tracks=Reference%2CMappability%20k%3D76%2CAnnotation%2CRB-1B%205%20days%20coverage%20reverse%2CRB-1B%205%20days%20introns%2CTentative%20reannotation%20RB-1B%2CCVI-988%205%20days%20coverage%20reverse%2CCVI-988%205%20days%20introns%2CTentative%20reannotation%20CVI-988&highlight= Panel D: https://mallorn.pirbright.ac.uk/browsers/MDV-annotation/?loc=GaHV2_MD5%3A105058..110282&tracks=Reference%2CMappability%20k%3D76%2CAnnotation%2CRB-1B%205%20days%20coverage%20reverse%2CRB-1B%205%20days%20introns%2CTentative%20reannotation%20RB-1B%2CCVI-988%205%20days%20coverage%20reverse%2CCVI-988%205%20days%20introns%2CTentative%20reannotation%20CVI-988&highlight=.

Our data confirms many MDV spliceforms previously reported in the literature. The splicing variant which was reported (20) between the ORF011 and ORF012 genes of MDV can be found in our RNASeq data – it is annotated on the genome browser as RB-1B-0865 or CVI-988-0883. In addition, we observed both spliced variants of pp38 in CEF cells infected with MDV strain RB-1B (RB-1B-0918 and RB-1B-0919 according to the nomenclature used in our reannotation) and one spliced variant of pp38 in cells infected with CVI-988 (CVI-988-0979). In the gC region, we observed the two previously described splicing transcripts (19): namely RB-1B-0874 and RB-1B-0875; or CVI-988-0920 and CVI-988-0921 in RB-1B and CVI-988 infected CEFs, respectively. Additionally, we observed three further novel spliced isoforms – RB-1B-0933, RB-1B-0876, and RB-1B-0877 in CEFs infected with the RB-1B strain; and CVI-988-1051, CVI-988-0922, and CVI-988-0923 in CEFs infected with strain CVI-988. We observe a higher diversity in transcripts of the ICP4 gene in cells infected with MDV strain CVI-988 than in cells infected with MDV strain RB-1B.

Although our data confirms many known spliceforms of MDV, we could not identify all of them. For example, we could not observe the spliceform between Meq and vIL8 loci which was described as meqΔC-BamL in MSB-1 cells or infected CEF cells (16, 23). Unlike other previous reports (17), we only observed one unspliced product for the Meq protein. In the IL8 region, we did not observe RLORF4/IL8 or RLORF5/vIL8 related transcripts (17). According to our data, an alternative start codon in vIL18 region could be employed to produce a novel transcript for vIL8 made of exons I and II of the gene, but representing an overlapping ORF (RB-1B-0928 or CVI-988-1019) to the main ORF of vIL8 (RB-1B-0936 or CVI-988-1068). In the vIL8 region, splicing between exon I and exon II would produce a shorter transcript than the full vIL8 transcript, with a length of 379 nucleotides and containing an immature stop codon at the end of the second splice site (RB-1B-0929 or CVI-988-1020).

### Selection of strain-dependent introns

By comparing the coverage in spliced reads at the same genomic positions between virulent (RB-1B) and vaccine (CVI-988) MDV strains, a list of viral introns showing the highest variation between strains was generated (Table 3, see Materials and Methods for a description of the procedure used). According to this coverage-based ranking and feasibility of real-time PCR probe design, three introns that were exclusively spliced in the CVI-988 transcriptome (indicated in Table 3 as I1, I2, and I3) were selected for further study. Introns I1 and I2 are highlighted in Figure 4, while intron I3 can be found in Figure 3.

### The kinetics of I1 expression in CEFs transfected with MDV strains CVI-988 and RB-1B

In order to determine the time window during which I1 was transcribed from the MDV genome, its time-dependent expression was subsequently determined using qRT-PCR on RNA isolated from CEFs infected with both MDV strains RB-1B and CVI-988. MDV is avidly cell associated, which makes it problematic to achieve synchronised expression of viral transcripts due to in vitro growth differences between attenuated CVI-988 and virulent RB-1B. Hence the time course experiment was performed by transfecting CEFs with equimolar amount of recombinant bacterial artificial chromosome (BAC) containing the complete genomes of MDV strains CVI-988 and RB-1B (24, 25). Expression levels were assessed from transfected cells harvested at nine different time points ranging from 0 to 90 hours post transfection, using GAPDH as a reference for relative quantitation (Figure 6 panel B).

**Figure 6.**
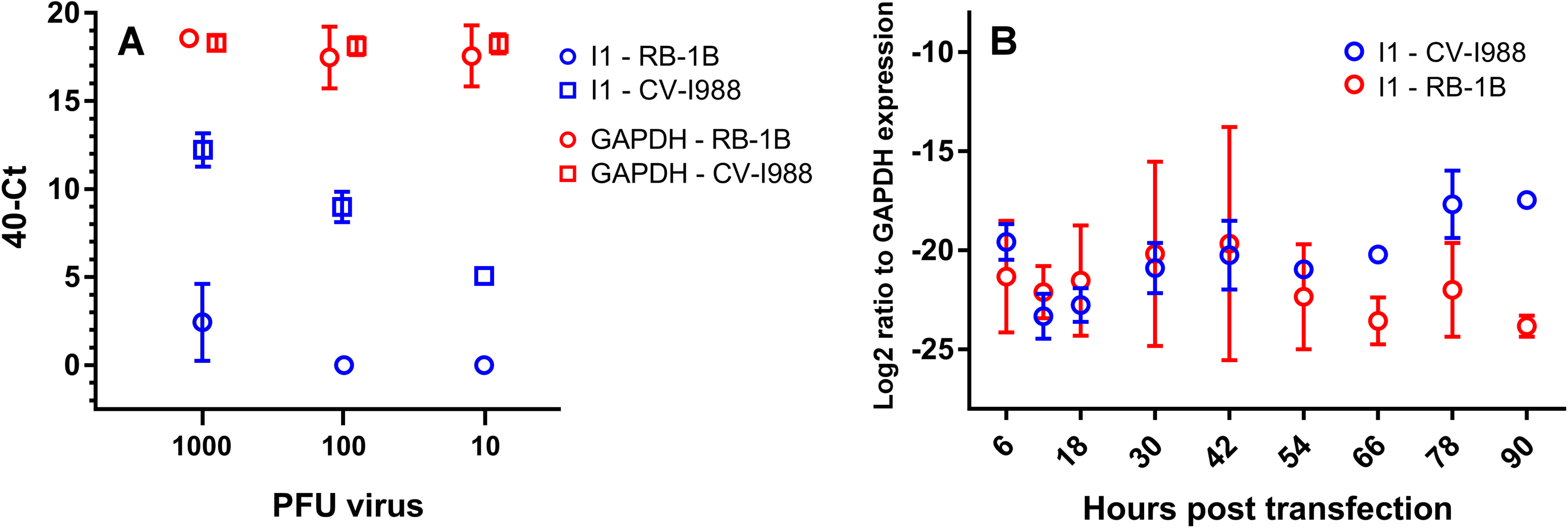
Panel A. Expression level of GAPDH and I1 at 5 days post infection in CEFs infected with 1000, 100 or 10 PFU of MDV strain CVI-988 or RB-1B. Three independent biological replicated were measured per PFU value. Note that the *x* scale is not continuous – there are only three, discrete, values corresponding to 1000, 100 and 10 PFU. Small horizontal deviations from such values have been introduced only to make the plot more readable. **Panel B**. **Expression of I1 in CEFs transfected with RB-1B or CVI-988 BAC DNA as a function of time.** Cells were harvested at time points 6, 12, 18, 30, 42, 54, 66, 78, and 90 hours post transfection. Expression levels of I1 were calculated and normalised to the corresponding levels for GAPDH (see Materials and Methods), and the ratios displayed on the graph for each time point. Three independent biological replicates were measured per time point. P-values are 0.37, 0.29, 0.50, 0.81, 0.88, 0.41, 0.008, 0.06, and 0.00067. P-values were computed with a multiple t-test.

Intron l1 is expressed at roughly equal levels in both MDV strains RB-1B and CVI-988 infected cells until 54 hours post transfection (the baseline being that GAPDH is about 4×10^6^ times more expressed than I1). However, starting from 66 hours post transfection the expression of I1 progressively increases (with differences becoming statistically significant) in cells transfected with MDV strain CVI-988. The peak is recorded at 90 hours post transfection, when the expression level is about 100 times greater in cells transfected with MDV strain CVI-988 than in cells transfected with MDV strain RB-1B. This finding agrees with the earlier observation deduced from RNA-sequencing data that at 3 days post infection, and increasingly so, I1 is expressed at exceedingly greater levels in CEFs infected by CVI-988 than in cells infected by RB-1B, making I1 a good candidate marker to discriminate between the two strains. A viral transcript was used to assess transfection efficiency, and its level was similar in CEF cells transfected with either of the strains (data not shown).

### I1 is detectable even at low viral loads

The expression differentials just described remain true at 5 days post infection across a wide range of virus infectious loads. The expression levels of I1 were quantified using relative RT-PCR with RNA isolated from 1.3×10^6^ CEF cells infected with 1,000, 100 or 10 PFU of MDV strains RB-1B or CVI-988 in 9 cm^2^ tissue culture dishes (Figure 6 panel A) – due to the cell-associated nature of MDV, we were unable to calculate MOI. The following lists the results:

1. The expression of the housekeeping gene, GAPDH remained unchanged at a 40-Ct value of ~18 across the different PFU levels (p = 0.48 based on a two-way repeated measures ANOVA test) and independent of the strain.
2. In CEFs infected by MDV strain CVI-988 the levels of expression of I1 also correlated very well with inoculum titres (Pearson’s r 0.98, p-value 6.1×10^−6^) and I1 was detectable across the whole PFU range.
3. I1 expression was undetectable in 1.3×10^6^ CEFs infected with 100 or 10 PFU of MDV strain RB-1B. In cells infected with 1,000 PFU I1 levels were still extremely low (averaged 40-Ct value 2.4).

Overall, our results support the use of intron I1 as a biomarker across a wide range of viral titres. Whenever it should be detectable – i.e. in CEFs infected by MDV strain CVI-988 starting from roughly 60 hours post infection – it remains so even at very low virus input, and its abundance at 5 days post infection in vitro shows a very clear and predictable linear relationship with virus input. We assumed that these findings would be generalizable to I2 and I3.

### Differential expression of I1, I2, and I3 in MDV strains RB-1B and CVI-988

At 5 days post infection all our three candidate introns (I1, I2, and I3) are differentially expressed in CEFs infected with MDV strains RB-1B and CVI-988. Such differential expression can be quantified using GAPDH as PCR calibrator.

Briefly, in order to precisely calculate expression levels, standard curves using PCR-amplified or cloned fragments of DNA representing I1, I2, I3, and GAPDH were first made and used to calculate PCR efficiency. Each amplicon was Sanger sequenced in order to confirm that the sequence identified by PCR corresponded with that of the splice junction predicted from RNA-sequencing data. The expression levels of the three biomarkers (I1, I2, I3) in CEFs infected by MDV strains RB-1B or CVI-988 at 5 days post infection (N=6) were computed and normalised against GAPDH, in order to achieve accurate calibration. Fresh virus stock obtained after only two passages was used for infection for both strains. Full details are described in Material and Methods.

The results are presented in Figure 7, and represent one of the main outcomes of this paper. When comparing expression in CEFs infected with MDV strain CVI-988 to expression in CEFs infected with MDV strain RB-1B, the greatest fold-change was observed for intron I1, which appeared to be expressed ~2800 times more when GAPDH was used as a calibrator. I2 and I3 exhibit fold-changes that are lower than those of I1 but still easily measurable and statistically significant (I2 vs. GAPDH: 6.3; I3 vs. GAPDH: 5.3) confirming the ranking originally deduced from RNA-sequencing data (Table 3).

**Figure 7.**
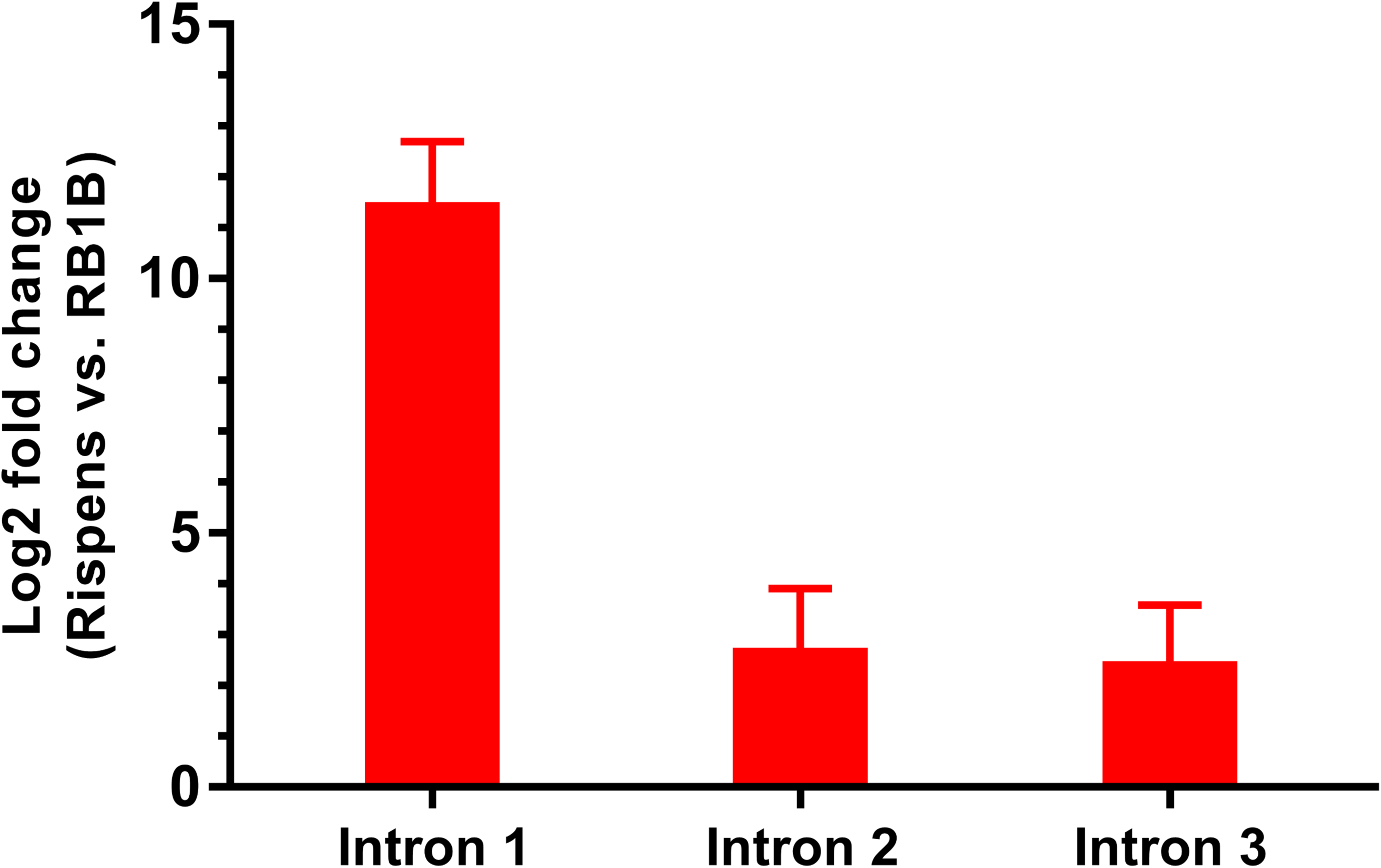
Differential expression in CEFs of introns I1-I3 in CVI-988 and RB-1B. Expression levels of introns I1, I2, and I3 were detected in CEFs infected with 500 PFU of MDV strains CVI-988 or RB-1B at 5 days post infection, and the ratio of the expressions in CVI-988 and RB-1B was computed for each intron (see labels on the x axis). Expression levels in RB-1B and CVI-988 were normalised against the expression level of GAPDH. Six independent biological replicates were performed.

## Discussion

The pathobiology of Marek’s disease virus is complex (26). This alphaherpesvirus is able to infect chicken cells and cause tumours in a wide variety of tissues. The outcome is that virions are shed into the environment as infectious dander (5), while cell free viruses can hardly been detected in vitro in the supernatants of any cell line. Some Mardivirus-specific genes have been identified and characterized, but more research is needed to understand the molecular mechanisms responsible for its complex life cycle, especially at the transcriptional level.

A number of studies based on high-throughput RNA-sequencing conducted with a number of technologies – including Illumina, Pacific Biosciences and Oxford Nanopore – have been published so far for all subfamilies of herpesviruses: the alphaherpesviruses herpes simplex virus (11, 12) and pseudorabies virus (27, 28); the betaherpesvirus cytomegalovirus (CMV); the gammaherpesvirus Epstein-Barr virus (EBV) and Kaposi’s sarcoma-associated herpesvirus (13, 14, 29). For several of those, the viral transcriptome has even been characterised for several tissues and biological conditions. A similar extensive characterisation for MDV has been lacking for a long time. In the past, some MDV transcripts had been determined through northern blot analysis and the nucleotide sequencing of a limited number of cDNA products, most notably the spliced variants of MDV’s oncogene Meq (16), vIL8 (17) and glycoprotein C (19). Very recently, a study of MDV infection in B cells has identified novel transcripts and genes (15). To expand on this, we report on the viral transcriptome of MDV using high-throughput short-read RNA sequencing on RNA isolated from CEF cells infected with MDV virulent strain – RB-1B (30) – and MDV attenuated vaccine strain – CVI-988 (31).

Interestingly, and consistent with what can be observed across the *Herpesviridae* family and in MDV-infected B cells, our results show that RNA splicing is pervasive in CEFs infected with these strains, giving rise to hundreds of so far unannotated spliceforms which biological significance remain undefined and in need of further investigation. The level of complexity we find is similar to what was observed for CMV (13) or EBV (29). For instance, in CMV infected cells a total of 751 ORF were identified which are transcribed and translated into proteins (13). During EBV reactivation, a high level of bidirectional transcription was observed (29). The complex transcription landscape that map to the UL region of MDV suggest that in addition to transcripts encoding early and late gene products other functional categories might be present, with a number of them possibly having regulatory functions in the control of gene products expression. This has been observed for both pseudorabies and herpesvirus simplex (12, 27, 28).

The intron I1 we identified could potentially represent a new protein-coding transcript which is transcribed from the UL49 locus of MDV strain CVI-988. It is reported that the VP22 protein of MDV is involved in cell cycle arrest, virus spread and histone association during MDV replication in vitro (32). Further studies will be required in order to investigate the involvement of the I1 transcript in the interaction between virus and host.

One should note that there are limitations to the sensitivity of the technique we employed in this study – in particular, spliceforms that are too weak with respect to the most abundant RNA species will not be detectable, depending on the number of sequencing reads generated during the experiment. In fact, as described in the Results section, a number of splicing isoforms identified in previous studies by qPCR could not be found in our data. Whether this is due to limited sensitivity or to the fact that such isoforms are specific to other experimental conditions – for instance, (23) used MSB1 cells rather than CEFs – the conclusion can only be that the potential size of the MDV transcriptome is even larger than what emerges from our study. That is confirmed by the novel transcripts identified in (15). Again, this is entirely similar to what can be seen for other herpesviruses – in previously studied CMV infected cells, the authors were unable to identify some of the previously annotated genes (13).

Further complementary investigations based on long-read technologies (e.g. Pacific Biosciences) will also be required to fully resolve the genomic structure of long spliced transcripts being made of many consecutive exons.

Even more interestingly, a significant number of viral splicing isoforms of MDV strains CVI-988 and RB-1B appear to be strain-specific, despite the high sequence similarity (>99%) between the two strains. We validated this finding by using a Taqman-based assay. The assay was proven to be reproducible. As far as we know such a striking result had not been previously reported in the literature, and it is not reported in (15) either. Not surprisingly, our preliminary analysis of the host transcriptome (data not shown) indicates that the splicing of host transcripts also depends on the virulence of the infected strains. We point out that the effect we notice cannot be ascribed to culture adaptation – as explained in the Material and Methods section, both MDV strainRB-1B and MDV strain CVI-988 stocks we used underwent only two passages before being used for infection. Likewise, what we observe cannot be explained away by the known fact that splicing tends to be highly unregulated in tissue culture cells for most herpesviruses, including MDV (17) – even in the presence of an equally permissive environment for splicing, MDV strain RB-1B appears to produce a much more limited number of splicing isoforms across the genome than MDV strain CVI-988. It is possible that the greater propensity for splicing exhibited by MDV strain CVI-988 indirectly derives from the fact that its attenuation was obtained by repeated passaging the strain in cell culture, but by now the trait has apparently been fixed in its genome, which is an interesting biological fact.

Taken together, our results suggest the possibility that different MDV strains might interact differently with the splicing machinery of the host; a different interaction of the spliceosome and associated proteins with the MDV genome in CEFs infected by the strain CVI-988 might result in a much richer splicing landscape. However, it is not possible at this stage to pinpoint the responsible for such differences at a molecular level – for instance, in the ICP27 region our data shows similar splicing patterns for MDV strain RB-1B and strain CVI-988 (with only one intron present in the strain CVI-988 and not in the strain RB-1B) and no differential expression (corrected p-value for differential expression computed by edgeR = 0.65), so it is unclear at the moment whether ICP27 might be responsible for what we observe.

As the results of RNA-sequencing describe the averaged collective nature of a relatively large number of infected cells, it is hard to say whether those differences are typical of every infected cell or just a smaller sub-population. In order to truly reveal the complexity of alternative splicing across diverse cell and tissue types, in future studies it might be prudent to sequence the RNA transcriptome of individual cells, especially from tissues (e.g. lymphocytes, tumours, FFE, etc.) of experimentally infected birds, and use proteomics to determine whether splice-variant transcripts are translated into protein products or just have RNA regulatory functions. We also plan to define the virus transcriptome in various tissues isolated from vaccinated/challenged bird that that are susceptible and resistant to Marek’s disease. That might help elucidate the molecular mechanisms underlying the diversity that we observe in splicing landscapes.

Overall, the data presented in this report suggest the presence of a large number of spliced viral transcripts in infected CEFs, allowing us to identify spliceforms that contain introns exclusive to the MDV strain CVI-988. Building upon such findings we have also designed primers and probes that can specifically detect such transcripts, thus effectively differentiating in CEFs the transcriptome of MDV strain CVI-988 from that of strain RB-1B. To the best of our knowledge, this study is the first one ever to propose a technique based on the differential detection of splicing events.

Obvious questions to be elucidated in the future are whether our results can be extended to other tissues/MDV strains; and whether differential expression of I1-I3 (or possibly other spliceforms yet to be identified) can also be detected in vivo.

## Acknowledgements

We thank Dr Luca Ferretti for providing statistical advice. The Pirbright Institute receives grant-aided support from the Biotechnology and Biological Sciences Research Council of the United Kingdom (projects BB/E/I/00007035, BB/E/I/00007036 and BBS/E/I/00007039). We also acknowledge support from BBSRC grants BBS/E/I/00001942 and BB/K011057/1. Mention of trade names or commercial products in this publication is solely for the purpose of providing specific information and does not imply recommendation or endorsement by the U.S. Department of Agriculture. This work was funded in part by USDA-ARS CRIS project number 6040-32000-074-00D.

## Authors’ contributions

The project was designed and conceived by ATA, PR and VN. Sample preparation for high-throughput sequencing was performed by ATA. Bioinformatics analysis was conducted by PR with assistance from SS. Real-time PCR experiments were designed and conducted by YS. The manuscript was prepared by PR, YS, SS and VN. All authors read and approved the manuscript.

## Materials and methods

### RNA-sequencing

#### Virus history

Splenocytes from a MDV strain RB1B infected bird (no.3345/KV16) were used to prepare a first passage of RB-1B virus stock. A second passage of MDV strain RB-1B was prepared using the first stock and used for RNA isolation, as described in the next section. MDV strain CV-I988 was prepared from CEF cells with two passage history after they were infected with Nobilis Rismavac vaccine virus. The passage 2 virus stock was used for RNA isolation, as described below.

#### Sample preparation

Confluent CEFs in a 75 cm^2^ flask (8.0 × 10^6^ cells in each flask) were infected with 1500 pfu of CVI-988 or RB-1B virus in DMEM containing 5% FBS and the antibiotic streptomycin (100μg/ml) and penicillin (100U/ml).

The flasks were incubated for five days at 37°C in 5% CO2 and RNA was isolated using Trizol (Ambion) according to the method described by the manufacturer. For each biological condition (CEFs infected with MDV RB-1B very virulent strain; and CEFs infected with MDV CVI-988 vaccine strain) two biological replicates were selected.

#### Sequencing

RNA samples were sequenced at the Centro Nacional de Análisis Genómico (Barcelona, Spain). Briefly, total RNA was assayed for quantity and quality using Qubit® RNA HS Assay (Life Technologies) and RNA 6000 Nano Assay on a Bioanalyzer 2100. The experimental protocol to construct stranded mRNA RNASeq libraries starting from the total RNA employed the TruSeq®Stranded mRNA LT Sample Prep Kit (Illumina Inc., Rev.E, October 2013). The initial input was 0.5 ug of total RNA for each sample. The size and quality of each final library were validated on an Agilent 2100 Bioanalyzer with the DNA 7500 assay (Agilent). Libraries were sequenced using TruSeq SBS Kit v3-HS in paired end mode with the read length 2×76bp. Each sample was sequenced in a fraction of a sequencing lane on a HiSeq2000/2500 machine (Illumina) following the manufacturer’s protocol, generating between 59 and 87 million paired end reads per sample. Images analysis, base calling and quality scoring of the run were performed using the manufacturer’s software Real Time Analysis (RTA 1.13.48) and were followed by generation of FASTQ sequence files with CASAVA 1.8. For this analysis the two biological replicates available for each MDV strain (RB-1B and CVI-988) were pooled together prior to analysis, in order to increase the overall read coverage.

### Bioinformatics selection of biomarkers

#### Primary analysis

Reads were subjected to preliminary quality control and processed with a workflow for primary data analysis based on the GEM mapper (33) which is an evolution of the method used to process the data produced by the GEUVADIS consortium (34). Contrary to other methods relying on accurate annotations, this one includes a highly sensitive de-novo intron detection step. This feature enabled accurate alignments in spite of errors or limitations in the available annotation of cellular transcripts, and an unbiased picture of splicing in different MDV strains.

It should be noted that separately aligning MDV RB-1B reads to the RB-1B genome and MDV CVI-988 reads to CVI-988 would have not been possible for this analysis, as during subsequent stages we needed to compare coverage of the same intron across different viruses. In order to be able to do so, reads from samples infected with different viruses (MDV strains RB-1B and CVI-988) were all aligned to the same MDV MD5 reference (NCBI accession number AF243438). Although in principle this procedure might decrease the amount of reads successfully mapped and potentially introduce artefacts, in practice it works well due to the high sequence similarity of the strains involved in the experiment – the conclusions presented in the paper are qualitatively identical when reads from both infections (RB-1B and CVI-988) are aligned to either MDV strains (MDV strain RB-1B, NCBI accession EF523390, or MDV strain CVI-988, accession DQ530348). As a relevant fraction of MDV genome is replicated twice – see for instance Figure 1, where mappability (35) for the genome is shown – reads aligning to more than one location of the genome were assigned to all locations, with normalisation 1/(number of alignments). Keeping only uniquely mapping reads, as is often done in RNA-seq data analysis, would have resulted in complete loss of signal in the repeated regions of the virus.

#### Tentative annotation of full spliced coding transcripts

The workflow used for primary analysis of each sample generated a list of introns, each one annotated with the number of spliced reads covering it. Introns having a coverage of 10 reads of more were kept, candidate exons were deduced from them, and an in-house script was used to generate all possible coding sequences of exons compatible with translation start and end signals present in MDV genome. Only putative transcripts leading to protein sequences longer than 35 amino acids were kept. This step was performed separately for the RB-1B and the CVI-988 data.

#### Selection of relevant MDV encoded introns

The intron list obtained after primary analysis was also split based on the following criteria: (i) viral introns that have a sufficient read coverage in CEFs infected with MDV strain RB-1B, and no coverage in CEFs infected with strain CVI-988; (ii) viral introns having sufficient read coverage in CEFs infected with strain CVI-988, and no coverage in CEFs infected with strain RB-1B. Introns were considered to be sufficiently populated whenever they had non-zero coverage in all biological replicates and the sum of their coverage across all replicates was ≥ 20 (arbitrary threshold). Finally, the list of introns was prioritised based on the geometric mean of the coverage in spliced reads across all replicates. An arbitrary final minimum threshold of 25 was used for the mean, in order to exclude from the list candidates with low level expression.

### Differential expression of transcripts

The differential expression for ICP27 mentioned in the Discussion was computed using edgeR (36) version 3.20.9.

### Data visualisation

The genome browser is based on a customised version of JBrowse (37).

### In-vitro validation of the internal RT-PCR control, GAPDH

#### Sample preparation

MDV strain RB-1B and CVI-988 were propagated as previously described (9, 10). To prepare samples for real time PCR, CEFs were infected in 9.8 cm^2^ wells (6 well plates, 1.3×10^6^ cells in each well) with 500 pfu of MDV strains RB-1B or CVI-988. Uninfected and infected cells were harvested for RNA purification using Trizol reagent (Ambion) as described by the manufacturer.

For the time course experiment, semi-confluent CEFs (80%) were transfected with 1 μg of RB-1B or CVI-988 BAC DNA clone using 10 μl of lipofectamine transfection reagent (Life Technologies) in 9.8 cm^2^ tissue culture plates. Three independent transfection was conducted to have 3 replicates (N=3). At 6, 12, 18, 30, 42, 54, 66, 87 and 90 hours post transfection, CEFs were harvested from the plates, washed with PBS, suspended in RLT buffer (Qiagen) and stored at −80°C until the time of RNA purification. RNA was isolated using RNeasy (Qiagen) and the contaminating DNA was destroyed by DNaseI treatment (New England Biolabs). cDNA was synthesized using RevertAid H minus reverse transcriptase (ThermoFisher Scientific) with random hexamers. Real-time PCR for I1 and GAPDH was conducted as described in (10, 38).

#### Primers and probes

Primers and probes were designed for each splicing isoforms using the PrimerQuest tool provided by Integrated DNA Technologies (IDT). Each probe was designed to span the splice junction, to avoid non-specific interaction between closely related splice variants. A list of primers and probes, sizes of the amplicons, and location of targeted introns is provided in Supplementary Table 3. GAPDH was used as the host gene. A pair of primers and a probe for the splice junction between exons 5 and 6 was designed to be used to detect the level of GAPDH cDNA. Each probe was labelled with 5’FAM reporter, ZEN™ and 3’-BHQ1 quenchers (Integrated DNA technology, IDT).

#### Relative quantitative RT-PCR

Real time qRT-PCR was performed using ABsolute blue qPCR mix (Thermo Scientific), primer pairs (each 0.4 uM) and probe (0.2 uM) for the splice junctions of interest. To generate a standard curve, splice junction amplicons were cloned in pGEM-T plasmid (Promega) and subjected to Sanger sequencing. Ten-fold serial dilutions were prepared to produce a range from 1 nM to 10 fM. Every real-time PCR run included, specific primers and probe for one of the splice junctions, the primers and probes for GAPDH, and the primers and probe for the virus control and were performed in triplicates. The reaction were processed on a 7500 Fast Real-time PCR system (Applied Biosystems) with reaction conditions specified by the master mix manufacturer. Data was collected and analysed using 7500 Software (v2.3, Applied Biosystems).

### Accession numbers

The raw sequencing reads have been deposited into the NCBI Sequencing Read Archive (SRA) under the project accession number PRJNA541962. The files corresponding to CEF cells infected with RB-1B and CVI-988 strains, each in two replicates, can be downloaded as accession numbers SRR9030404, SRR9030405, SRR9030406, and SRR9030407.

